# MultiPaths: a Python framework for analyzing multi-layer biological networks using diffusion algorithms

**DOI:** 10.1101/2020.08.12.243766

**Authors:** Josep Marín-Llaó, Sarah Mubeen, Alexandre Perera-Lluna, Martin Hofmann-Apitius, Sergio Picart-Armada, Daniel Domingo-Fernández

## Abstract

**Summary:** High-throughput screening yields vast amounts of biological data which can be highly challenging to interpret. In response, knowledge-driven approaches emerged as possible solutions to analyze large datasets by leveraging prior knowledge of biomolecular interactions represented in the form of biological networks. Nonetheless, given their size and complexity, their manual investigation quickly becomes impractical. Thus, computational approaches, such as diffusion algorithms, are often employed to interpret and contextualize the results of high-throughput experiments. Here, we present MultiPaths, a framework consisting of two independent Python packages for network analysis. While the first package, DiffuPy, comprises numerous state-of-the-art diffusion algorithms applicable to any generic network, the second, DiffuPath, enables the application of these algorithms on multi-layer biological networks. To facilitate its usability, the framework includes a command line interface, reproducible examples, and documentation. To demonstrate the framework, we conducted several diffusion experiments on three independent multi*-omics* datasets over disparate networks generated from pathway databases, thus, highlighting the ability of multi-layer networks to integrate multiple modalities. Finally, the results of these experiments demonstrate how the generation of harmonized networks from disparate databases can improve predictive performance with respect to individual resources.

**Availability:** DiffuPy and DiffuPath are publicly available under the Apache License 2.0 at https://github.com/multipaths.

## 1. Introduction

Emergent properties of biological processes primarily arise from complex interactions linking physical entities which, in turn, can build up complex biological networks, such as metabolic, signalling and regulatory. The use of these networks has become commonplace for a variety of analytic tasks, yet integrated networks have been shown to be more robust resources for analytic usage (Huang *et al.* 2018). Thus, several frameworks, such as Bio2RDF (Belleau *et al.* 2008), have been proposed to facilitate the integration of these networks from heterogeneous sources.

Numerous methods for network analysis derived from graph theory have been adapted for a broad range of applications in the biomedical domain including target prioritization, gene prediction and patient stratification (Cowen *et al.,* 2017). Amongst these methods, network propagation or diffusion, in particular, comprises a broad family of algorithms that infer node labels based on the sharing of labels through network connections.

Though a wide variety of algorithms exist, user-friendly software that can enable researchers to implement and compare several methods are lacking. Not only does this impede their adoption and reproducibility, but it also compels researchers to re-implement the algorithms for their particular needs. While recently, the R packages diffuStats and RANKS (Picart-Armada et al., 2017; Valentini et al., 2016) have addressed this issue, a framework that offers a pipeline to build harmonized networks from biological databases along with an array of ready-to-use diffusion algorithms has yet to be established.

Here, we present MultiPaths, a Python framework for the analysis of multi-*omics* data by various diffusion algorithms on harmonized networks from custom or predefined selections of biological databases. We demonstrate how MultiPaths enables contextualizing multi*-omics* experiments by presenting an application scenario on multiple datasets containing transcriptomics, metabolomics, and miRNomics data.

## 2. Implementation

The MultiPaths framework contains two independent Python packages: DiffuPy and DiffuPath. While DiffuPy is specifically designed for the implementation of diffusion algorithms, DiffuPath is capable of both generating harmonized biological networks, and running the algorithms over these networks. Their functionalities can be accessed programmatically and via a command line interface (CLI) for non-bioinformaticians. Their modular design eases the inclusion of network resources and algorithms in future releases.

### 2.1. DiffuPy

The first of the two packages in the framework, DiffuPy, enables propagating user-defined labels, either as lists of entities or lists of entities with their corresponding quantitative values, on a user-defined network. DiffuPy comprises four diffusion scores and five graph kernels that can be run on generic networks on different formats **(Supplementary Text)**.

### 2.2. DiffuPath

The second package, DiffuPath, wraps the generic diffusion algorithms from DiffuPy and applies them to biological networks. To that end, DiffuPath comprises a comprehensive pipeline that extends from the generation of harmonized networks from multiple biological databases to the visualization and analysis of the diffusion results **(Supplementary Text)**. The pipeline provides a user-friendly CLI that enables users to create customized networks from a pool of databases or predefined collections, directly run diffusion algorithms on these networks, and analyze them in a few commands.

### 3. Application

To demonstrate the framework, we run various diffusion algorithms from DiffuPy on four networks corresponding to four pathway databases generated through DiffuPath. The input labels for the diffusion derive from three independent datasets containing differential entities from three *-omics* modalities: transcriptomics, metabolomics, and miRNomics. The four networks consist of three well-established pathway databases: KEGG, Reactome, and WikiPathways (Kanehisa et al., 2016; Fabregat et al., 2017; Slenter et al., 2017) as well as their combined representation, PathMe (Domingo-Fernández *et al.,* 2019). Our hypothesis is that by integrating the three resources, PathMe covers a larger scale of interactions and entities as well as a broader range of interaction and modality types which can ultimately serve to improve prediction performance.

For each of the three datasets which investigated specific biological processes, we compared the prediction performance of the various diffusion algorithms in identifying genes, metabolites and miRNAs relevant to that particular biological process. This was repeated for each of the four networks and the performance was evaluated using a repeated holdout approach. For the raw diffusion scores, the distribution of area under the ROC curve (AUROC) scores indicated a significant improvement in prediction performance of the integrated multi-layer network over each of the individual databases **(Figure 1)**.

**Figure 1.**
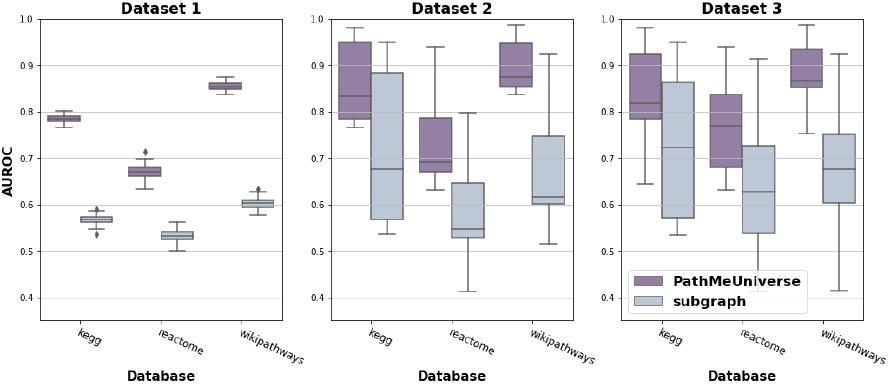
Prediction performance of raw diffusion over the integrated PathMe network and the correspondent database subgraph for three multi-*omics* datasets. Each box plot shows the distribution of the area under the ROC curves (AUROCs) over 100 repeated holdout validations. Details on the evaluation can be found in the **Supplementary Text.**

## 4. Discussion

This work has presented the first Python framework that implements numerous diffusion algorithms along with a pipeline to build customized harmonized networks from multiple biological databases. The importance of this integration is highlighted by our three case scenarios where a harmonized network leverages three *-omics* modalities (Di Nanni *et al.,* 2020) to increase predictivity in line with Huang *et al.* (2018). Furthermore, the integrated networks contain additional entities like biological processes and clinical readouts (e.g., symptoms and diseases), allowing a rich contextualisation of the experimental readouts **(Supplementary Text)**. This case scenario demonstrates the utility of diffusion algorithms to provide the interpretation of biological networks in the context of pathways, and thereby, elucidate the properties of biological processes underlying these networks. Additionally, the generalization and flexibility of the software also enables users to conduct analyses from biological networks to generic networks from other fields, such as social media. As a final remark, although large-scale and integrated multi-layer networks can improve prediction performance, greater computational power is required as the size of a network grows.

## Additional File

### Outline

1. DiffuPy
2. DiffuPath
3. Software Design
4. Case Scenario

### Supplementary Text

#### 1. DiffuPy

DiffuPy is a Python package that has been designed to run network propagation algorithms on generic networks (**Supplementary Figure 1)**. DiffuPy implements four existing network propagation algorithms and five graph kernels, introducing them to the Python community as, until now, they were limited to other programming languages such as R (e.g., diffuStats (Picart-Armada *et al.,* 2017a) and RANKS (Valentini *et al.,* 2016)). In its implementation, DiffuPy leverages NetworkX (Hagberg *et al.,* 2008), one of the most commonly used packages for working with networks in Python.

Scores including **raw**, **ml** and **gm** are computed as *f* = *Ky*, where *f* is the score vector, *K* is the graph kernel and *y* contains the input scores. These scores differ on how *y* is codified when the input is defined as a set of positive, negative and unlabelled nodes **(Supplementary Table 1)**. A fourth score, **z**, corrects node-wise **raw** scores with an exact z-score under the permutation of labelled nodes. **z** has been found to be better suited over **raw** in the interpretation of metabolomics data using heterogeneous knowledge graphs from the KEGG database (Picart-Armada *et al.* 2017b). On the other hand, several options are available for choosing *K* **(Supplementary Table 2)**, which dictates how the propagation behaves. By default, the regularised Laplacian kernel is used, which is a commonly used fluid or heat propagation model (Cowen *et al.,* 2017).

Furthermore, DiffuPy provides a command line interface (CLI) for improving its usability such that analyses can be conducted on the terminal, as per the guidelines in Grüning *et al.* (2019). The most recent version of the software allows users to input several standard network formats such as SIF or GraphML, facilitating its usage across different formats. This in turn also enables running the diffusion algorithms on non-biological networks from other fields, such as social media. Finally, the source code of the DiffuPy Python package is available at https://github.com/multipaths/diffupy, its latest documentation can be found at https://diffupy.readthedocs.io and its distributions can be found on PyPI at https://pypi.org/project/diffupy.

**Supplementary Table 1.**
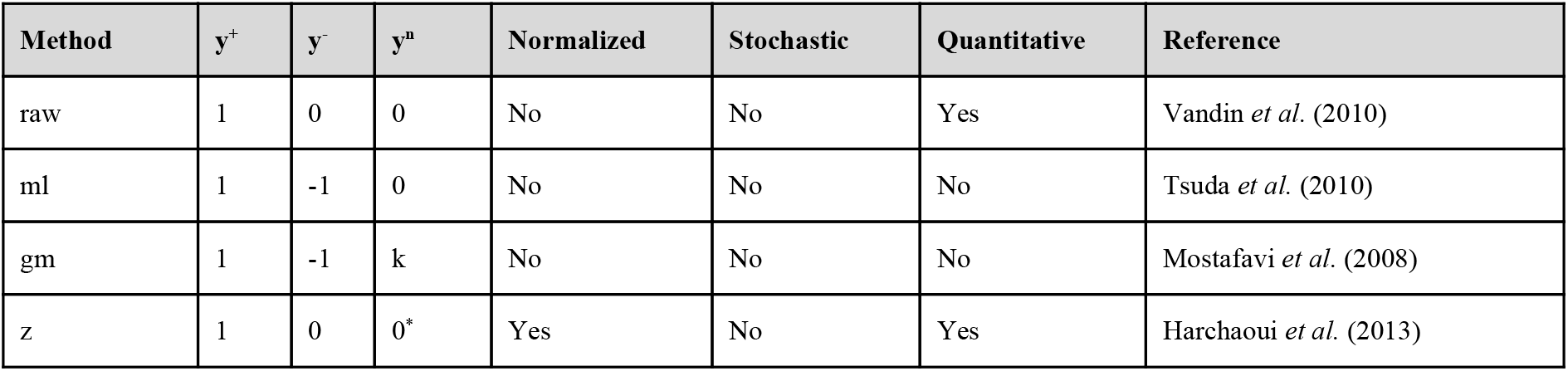
Diffusion methods implemented in DiffuPy. Methods differ on how the input labels are codified in the propagation (positive entities y^+^, negatives y^-^ and unlabelled y^u^). Besides the positive/negative/unlabelled paradigm, **raw** and **z** also accept quantitative inputs, which are passed to the propagation directly. The **z** method offers a statistical normalization, specifically an exact z-score, based on the permutation of the labelled nodes.

| Method | $y^+$ | $y^-$ | $y^u$ | Normalized | Stochastic | Quantitative | Reference |
| --- | --- | --- | --- | --- | --- | --- | --- |
| raw | 1 | 0 | 0 | No | No | Yes | Vandin *et al.* (2010) |
| ml | 1 | -1 | 0 | No | No | No | Tsuda *et al.* (2010) |
| gm | 1 | -1 | k | No | No | No | Mostafavi *et al.* (2008) |
| z | 1 | 0 | 0* | Yes | No | Yes | Harchaoui *et al.* (2013) |

**Supplementary Table 2.** Graph kernels implemented in DiffuPy. The regularisation function is the transformation applied to the graph Laplacian spectrum, as defined in Smola (2003), that characterises each kernel. σ^2^ > 0, *a* ≥ 2 and *p* ≥ 1 are kernel parameters.

| Kernel | Regularisation function | Reference |
| --- | --- | --- |
| Regularised Laplacian | $r(\lambda) = 1 + \sigma^2 \lambda$ | Smola *et al.* (2003) |
| Diffusion process | $r(\lambda) = \exp(\frac{\sigma^2}{2} \lambda)$ | Smola *et al.* (2003) |
| p-Step random walk | $r(\lambda) = (a - \lambda)^{-p}$ | Smola *et al.* (2003) |
| Inverse cosine | $r(\lambda) = (\cos(\lambda \frac{\pi}{4}))^{-1}$ | Smola *et al.* (2003) |
| Commute-time kernel | $r(\lambda) = \lambda$ | Yen *et al.* (2007) |

#### 2. DiffuPath

The primary goal of DiffuPath is to act as a connector between harmonized biological networks and the array of diffusion algorithms implemented in DiffuPy. DiffuPath leverages two resources, PathMe (Domingo-Fernández *et al.,* 2019) and Bio2BEL (Hoyt *et al.,* 2019), by integrating biological databases into a common network-like schema (i.e., Biological Expression Language; BEL) to offer a pipeline that can generate harmonized networks based on the database selection from the user **(Supplementary Figure 1)**. In total, users can select from nine single-resource databases **(Supplementary Table 3)**. By using the harmonized network of all available databases, one could cover up to three *-omics* modalities (i.e., genomics/proteomics, miRNAs, metabolomics) as well as clinical endpoints (i.e., symptoms, side effects), biological processes (i.e., pathways) and diseases. To facilitate users choosing a selection, we also provide predefined collections of databases based on the goal of the study and dataset characteristics **(Supplementary Table 4)**. Furthermore, DiffuPath contains functions to explore the overlap between the input and the network in the *views* module. Finally, the source code is available at https://github.com/multipaths/diffupath, its latest documentation can be found at https://diffupath.readthedocs.io, and its distributions can be found on PyPI at https://pypi.org/proiect/diffupath.

**Supplementary Figure 1.**
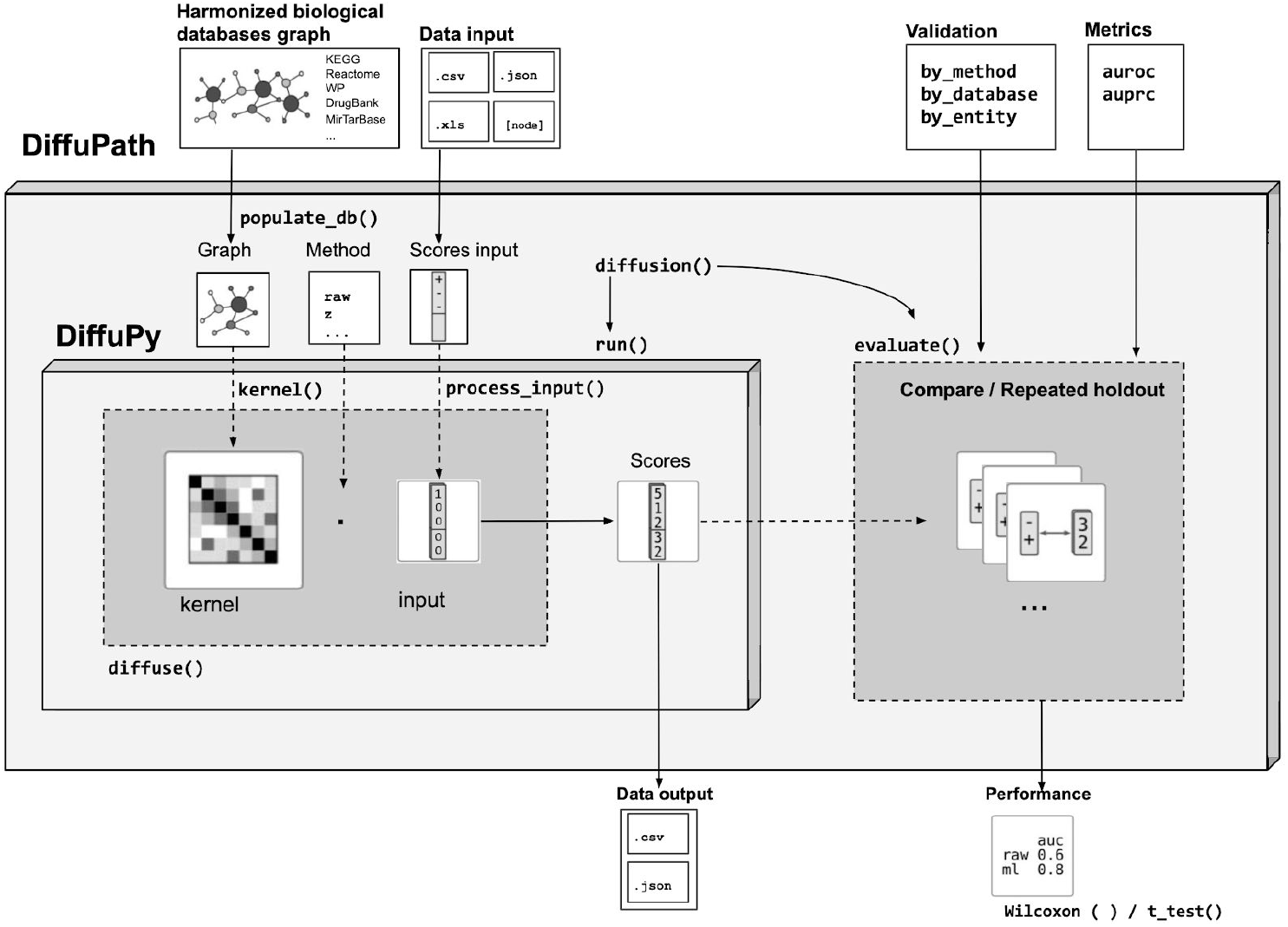
DiffuPath and DiffuPy framework.

**Supplementary Table 3.** List of biological databases available from DiffuPath.

| Name | Description | Reference |
| --- | --- | --- |
| DDR | Disease-disease associations | Menche *et al.* (2015) |
| DrugBank | Interactions between drugs and drug targets with over 10,000 drugs | Wishart *et al.* (2018) |
| Gene Ontology (GO) | Flexible hierarchy of tens of thousands of biological processes | Ashburner *et al.* (2000) |
| HSDN | Associations between thousands of diseases with hundreds of symptoms | Zhou *et al.* (2014) |
| KEGG | Multi-omics interactions present in hundreds of biological pathways | Kanehisa *et al.* (2016) |
| MirTarBase | Nearly 300,000 experimentally validated interactions between miRNA and their targets | Huang *et al.* (2020) |
| Reactome | Multi-omics interactions present in thousands of biological pathways | Fabregat *et al.* (2018) |
| SIDER | Associations between over a thousand drugs and side effects | Kuhn *et al.* (2016) |
| WikiPathways | Multi-omics interactions present in hundreds of biological pathways | Slenter *et al.* (2017) |

**Supplementary Table 4.** Predefined collections available in DiffuPath.

| Collection | Databases | Description |
| --- | --- | --- |
| #1 | KEGG, Reactome, and WikiPathways | -omics and biological processes/pathways |
| #2 | KEGG, Reactome, WikiPathways, and DrugBank | -omics and biological processes/pathways with a strong focus on drug/chemical interactions |
| #3 | KEGG, Reactome, WikiPathways, and MirTarBase | -omics and biological processes/pathways enriched with miRNAs |

**Supplementary Table 5.**
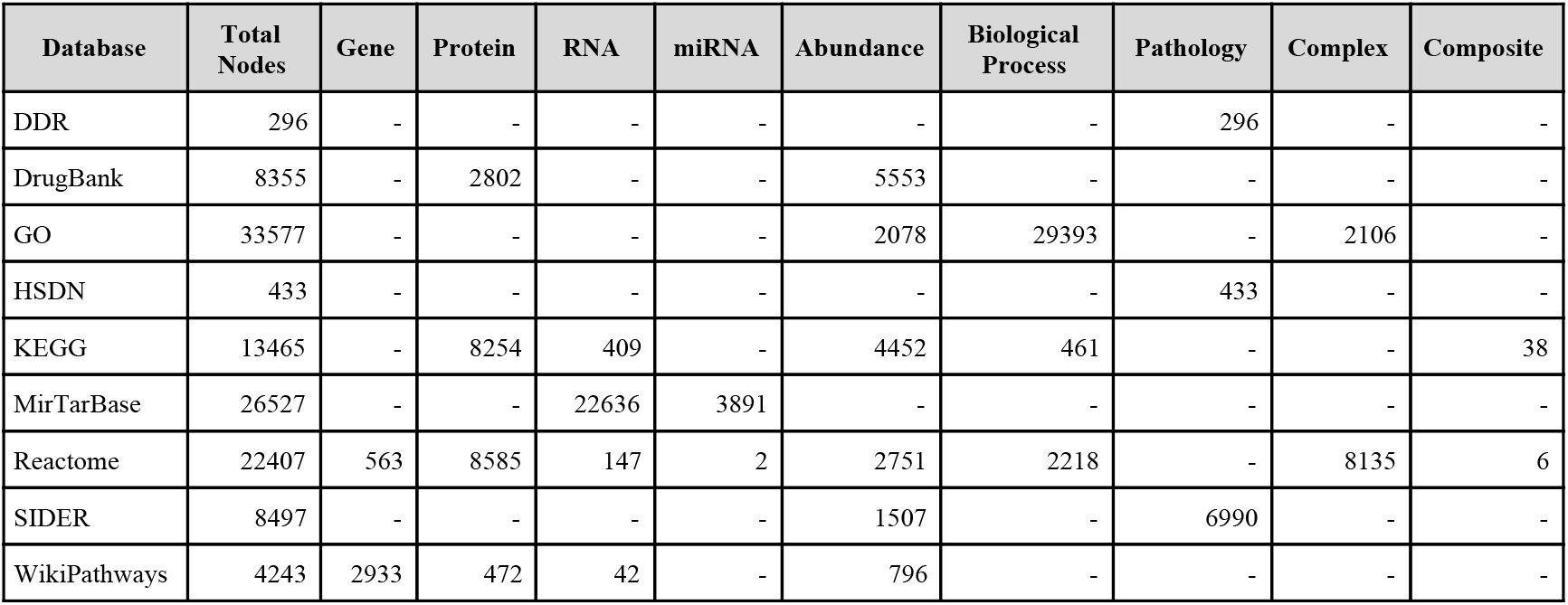
Statistics of the entity types for each of the individual databases available in DiffuPath.

**Supplementary Table 6.**
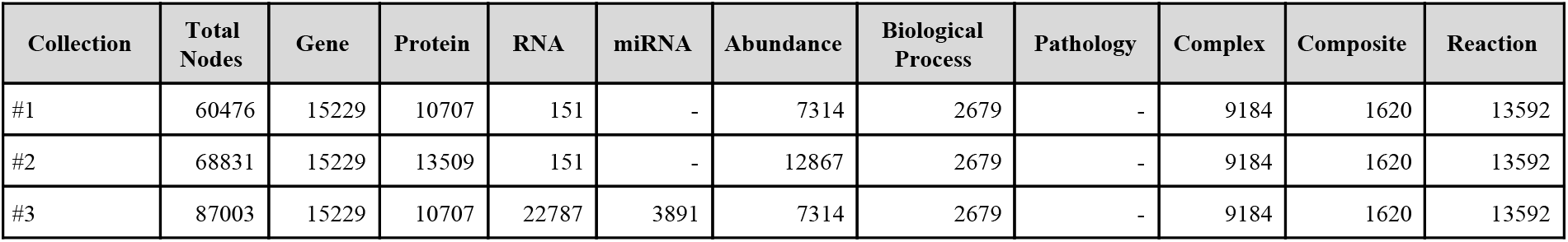
Statistics of the entity types for each of the collections available in DiffuPath.

#### 3. Software Design

Both DiffuPy and DiffuPath have a tool chain consisting of pytest (https://github.com/pytest-dev/pytest) as a testing framework, coverage (https://github.com/nedbat/coveragepy) to assess testing coverage, sphinx (https://github.com/sphinx-doc/sphinx) to build documentation, flake8 (https://github.com/PyCQA/flake8) to enforce code and documentation quality, setuptools (https://github.com/pypa/setuptools) to build distributions, pyroma (https://github.com/regebro/pyroma) to enforce package metadata standards and tox (https://github.com/tox-dev/tox) as a build tool to facilitate the usage of each of these tools in a reproducible way. These packages leverage community and open source resources to improve their usability by using Travis-CI (https://travis-ci.com) as a continuous integration service, monitoring testing coverage with Codecov (https://codecov.io) and hosting its documentation on Read the Docs (https://readthedocs.org).

#### 4. Case Scenario

##### 4.1. Network

The harmonized network chosen for the case scenario comprises three major pathway databases (i.e., KEGG; Kanehisa *et al.,* 2016, Reactome; Fabregat *et al.,* 2018, and WikiPathways; Slenter *et al.,* 2017). This integrated network was selected as it comprises pathway information from three highly-cited and open-sourced databases, covers a broad spectrum of human pathways, and incorporates three *-omics* modalities, thus enabling enhanced predictive power. The harmonized network of these three resources was generated using the CLI of DiffuPath leveraging its close integration with PathMe (Domingo-Fernández *et al.,* 2019). Because DiffuPath permits the separation of modalities, diffusion scores can be stratified by node type. For the case scenario, we thus selected PathMe as it incorporates multiple entity types (e.g., genes, miRNAs, and metabolites) that are also found in the multi-*omics* datasets chosen. Although edges can have attributes and directionality, the four networks used for label propagation were regarded as simple graphs (i.e., undirected, unweighted, no multi-edges or loops).

The statistics, including the number and type of nodes and edges of individual pathway database networks that DiffuPath provides are displayed in **Supplementary Table 5.** Group nodes, such as protein complexes and protein families, were represented through a group node formalism in which groups of entities were denoted as individual nodes and were connected to their neighbours by single edges. By selecting this representation approach, we sought to preserve biological context. Furthermore, we performed two additional preprocessing steps for the PathMe network. Firstly, isolated nodes were removed from the network. Secondly, if any proteins, RNAs and/or genes were equivalent, these nodes were collapsed into a single node (i.e., gene). The difference in statistics between collapsed and non-collapsed nodes are presented in **Supplementary Table 6 and Supplementary Figure 2**, respectively.

**Supplementary Figure 2.**
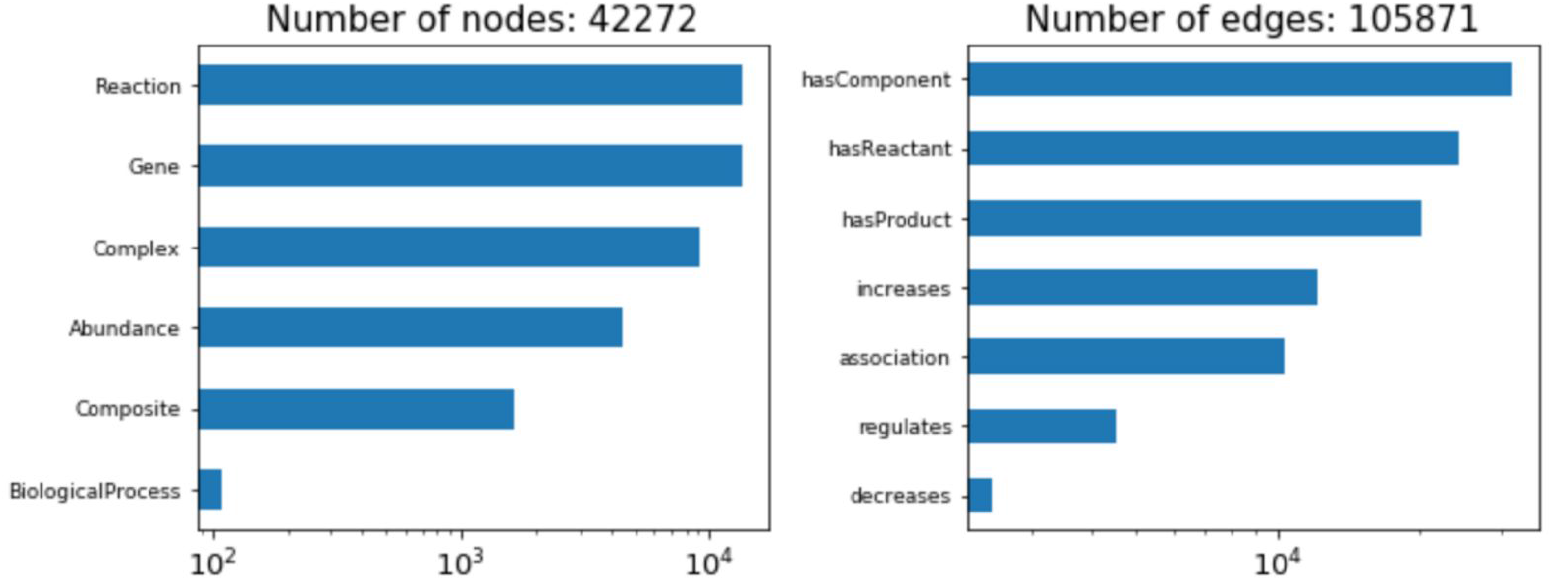
Node and edge statistics of the PathMe network.

##### 4.2. Datasets

In selecting the multi-*omics* datasets used in the case scenario for DiffuPath validation, the following criteria were applied:

- Each dataset had to contain three *-omics* modalities: molecular readouts on genes, metabolites and miRNAs.
- Each dataset had to be derived from human experiments.
- The original publication had to include differential analysis in order to avoid any bias in the preprocessing steps as well as include biological interpretation for validation purposes.

Following these criteria, three datasets were obtained and each was downloaded and customly parsed using DiffuPy processing tools. Data from the datasets was then mapped to the network, details of which are given below.

The **first dataset** comprises multi *-omics* experiments aimed to investigate the effect of exposure to cyclosporin A in hepatic cells (Van den Hof *et al.,* 2015). The authors quantified gene and miRNA expression as well as metabolomics and integrated the three *-omics* modalities to conduct an integrated pathway analysis. **Supplementary Figure 3** shows the overlap of each of these three modalities while **Supplementary Figure 4** depicts distributions of input measurements by entity type.

**Supplementary Figure 3.**
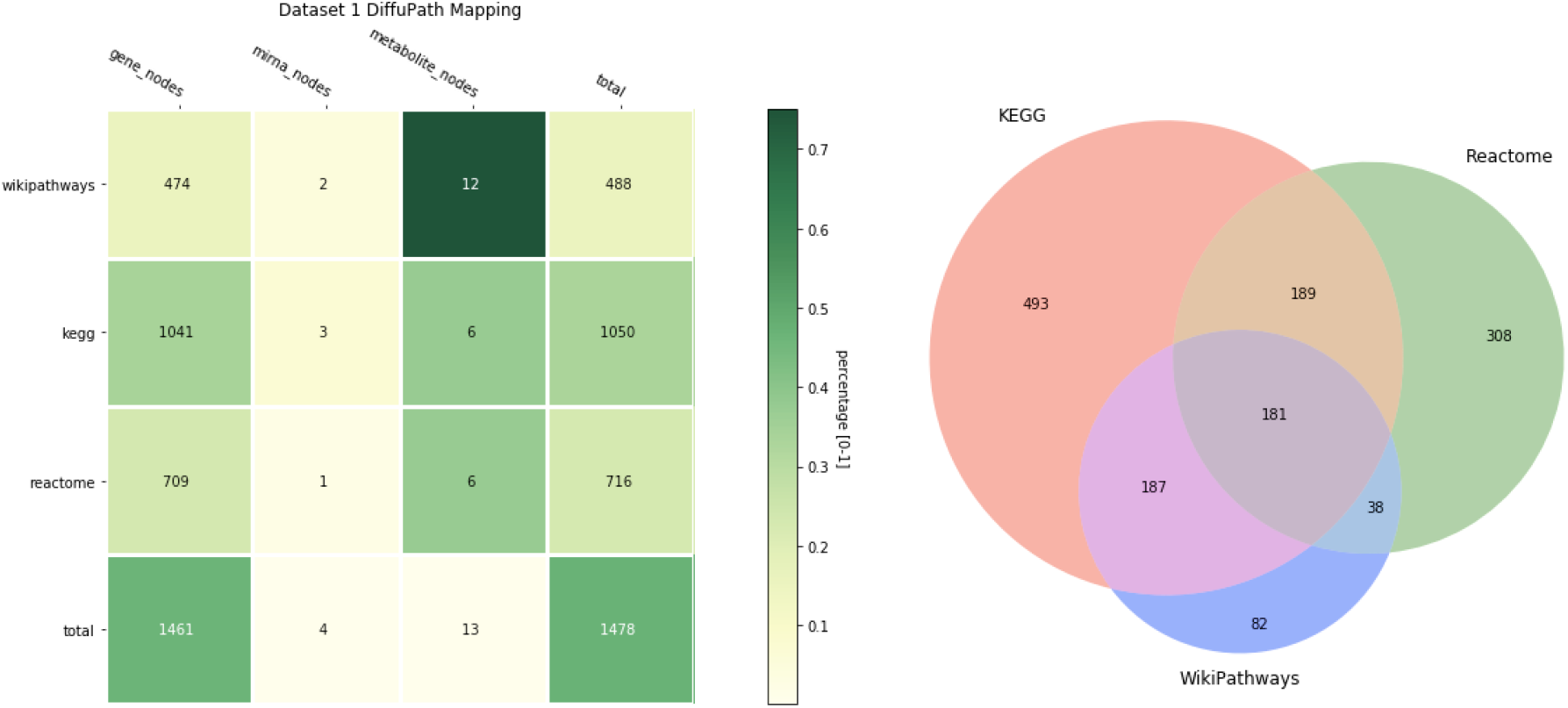
Mapping between dataset 1 and the network, stratified by entity modality (x axis) and by database (y axis) (left). The Venn diagram explores the overlap between multiple databases for the mapped entities (right). Colours in each cell of the heatmap represent the percent coverage of each modality for each database, except for the last row, where the denominator is the total amount of original entities in the dataset. The number in each cell corresponds to the total number of entities of a specific modality after mapping the input. The final column of the heatmap denotes the total number of entities in each of the three databases and the bottom row corresponds to the sum total of each modality across databases. Stratification by modality was performed in order to prevent bias of the results towards the most frequently occurring entity.

**Supplementary Figure 4.**
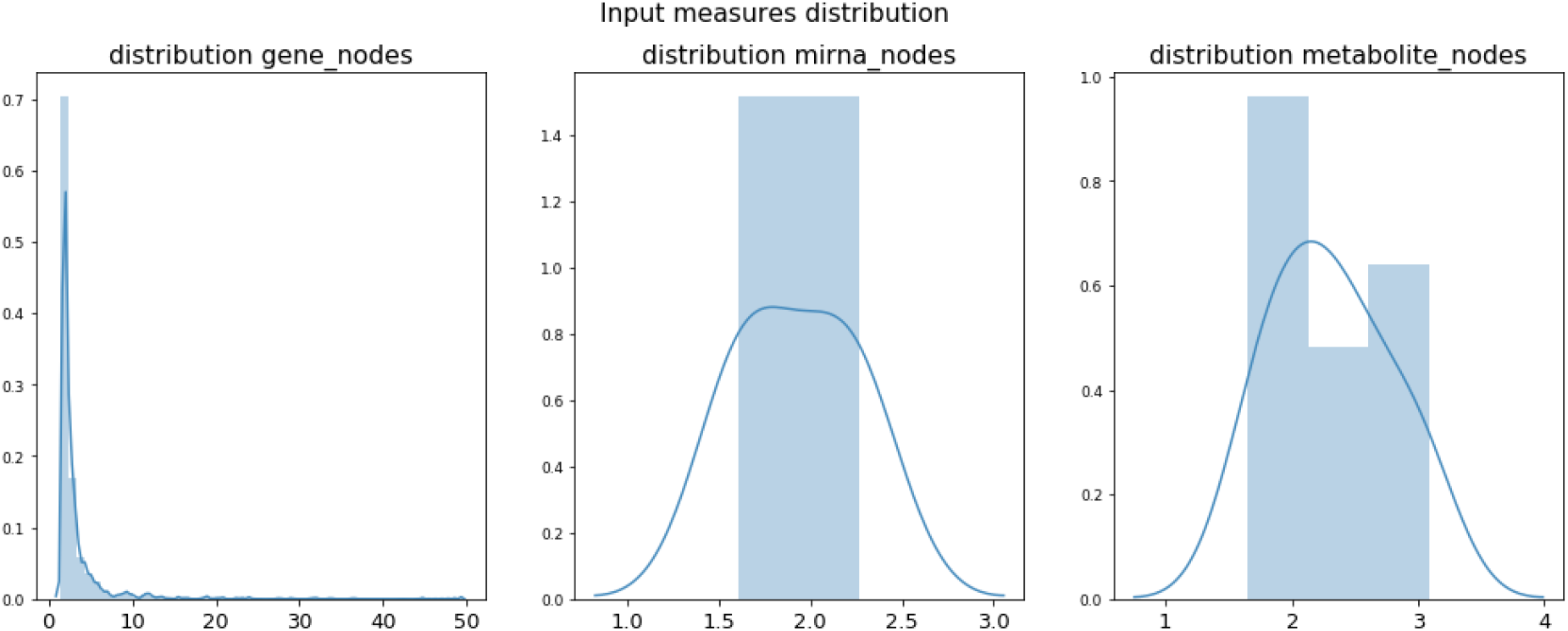
Distribution of the selected input measurements (i.e., absolute log_2_ fold changes) for dataset 1 stratified by *-omics* modality. Negative and positive raw values have been normalized by calculating their absolute value since the results of the diffusion need to be sorted in order to calculate the AUC values. The distribution of genes is relatively dispersed compared to the uniform distribution of miRNAs and metabolites. This can be attributed to the successful mapping of a larger proportion of genes than miRNAs or metabolites, thus resulting in a larger variance.

The **second dataset** aimed at identifying major metabolic pathways and potential biomarkers involved in prostate cancer (Ren *et al.,* 2016). Relatively fewer measurements of gene expression and metabolites were taken for this dataset, from which pathway information was inferred. **Supplementary Figure 5** shows the overlap of each of these two modalities

**Supplementary Figure 5.**
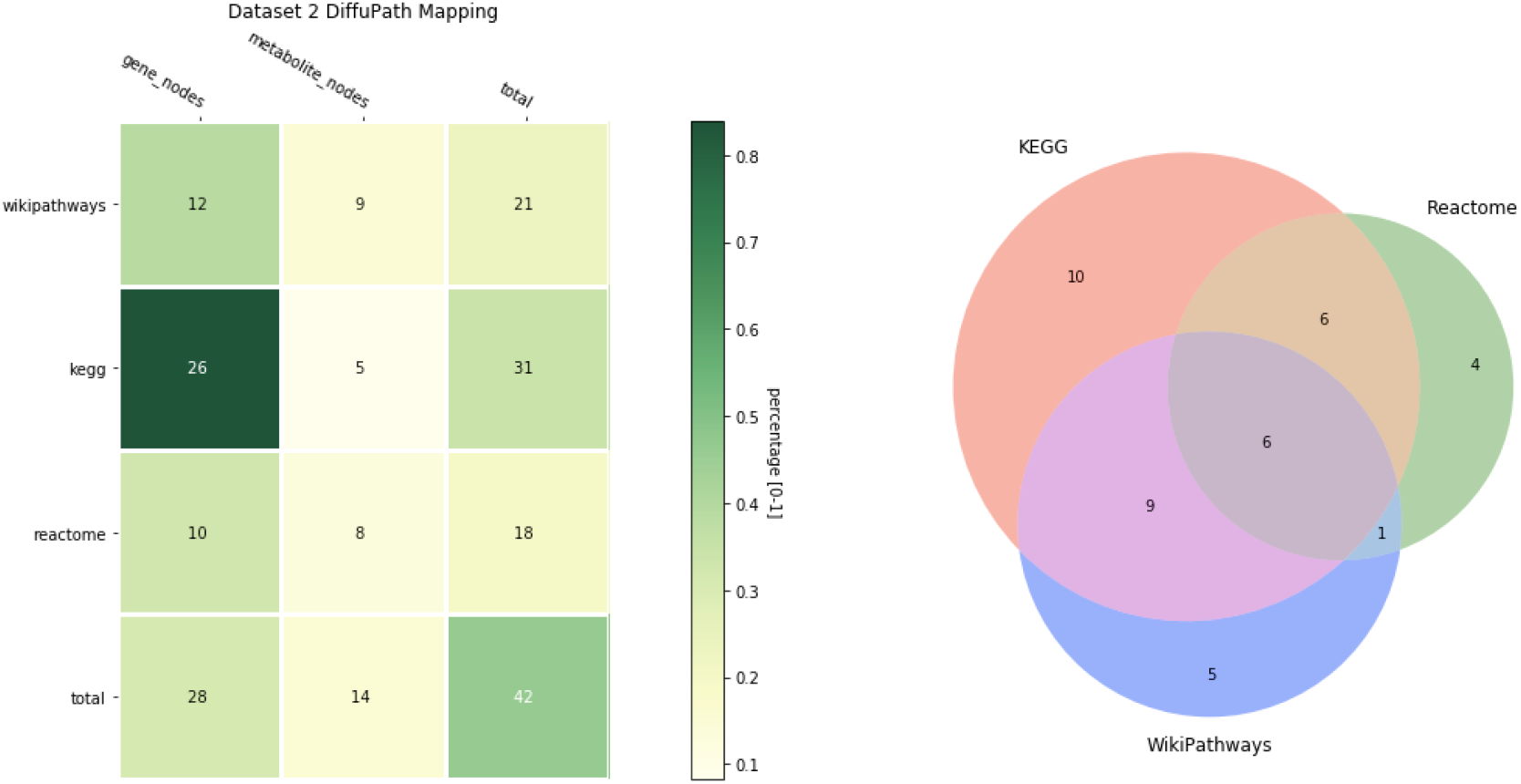
Mapping between dataset 2 and the network, stratified by entity modality (x axis) and by database (y axis) (left). The Venn diagram explores the overlap between the databases for the mapped entities (right). Colours in each cell of the heatmap represent the percent coverage of each modality for each database, except for the last row, where the denominator is the total amount of original entities in the dataset. The number in each cell corresponds to the total number of entities of a specific modality after mapping the input. The final column of the heatmap denotes the total number of entities in each of the three databases and the bottom row corresponds to the sum total of each modality across databases, Stratification by modality was performed in order to prevent bias of the results towards the most frequently occurring entity.

The **third dataset** investigates the carcinogenic mechanisms that differentiate intrahepatic cholangiocarcinoma from hepatocellular carcinoma (Murakami *et al.,* 2015). This dataset contains several genes, metabolites and one miRNA, as shown in **Supplementary Figure 7.**

**Supplementary Figure 6.**
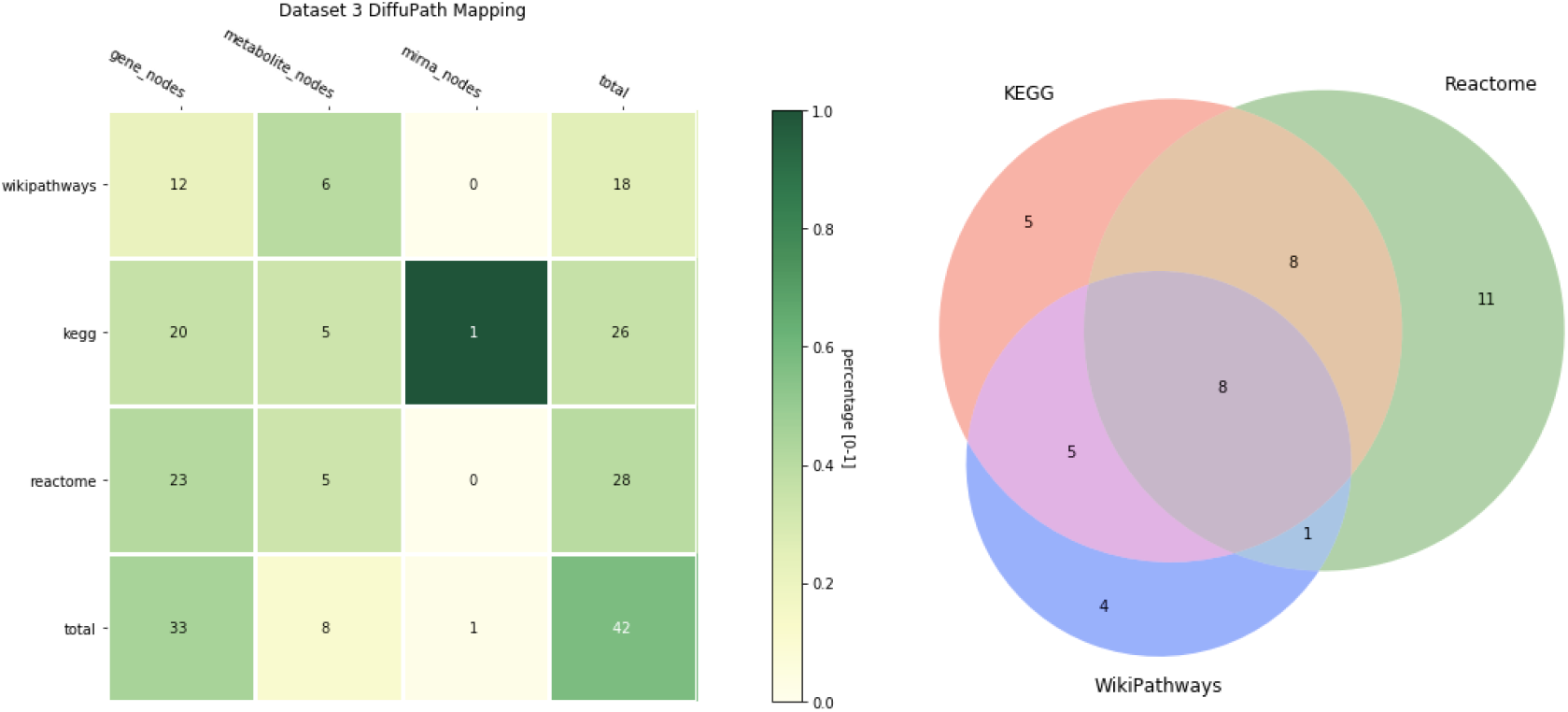
Mapping between dataset 3 and the network, stratified by entity modality (x axis) and by database (y axis) (left). The Venn diagram explores the overlap between the databases for the mapped entities (right). Colours in each cell of the heatmap represent the percent coverage of each modality for each database, except for the last row, where the denominator is the total amount of original entities in the dataset. The number in each cell corresponds to the total number of entities of a specific modality after mapping the input. The final column of the heatmap denotes the total number of entities in each of the three databases and the bottom row corresponds to the sum total of each modality across databases, Stratification by modality was performed in order to prevent bias of the results towards the most frequently occurring entity.

##### 4.3. Diffusion evaluation specifications

Network propagation has been previously applied to knowledge graphs and has been shown to benefit from statistical normalization in pathway analysis on heterogeneous networks. Raw and normalized z diffusion scores implemented in diffuStats (Picart-Armada *et al.,* 2017a) were adapted into DiffuPy, giving the user the flexibility to select the statistical background of the analysis (choosing between z normalization or other diffusion procedures lacking statistical normalization). Raw and z normalization label diffusion were compared (see “evaluation by method”) together with two baselines: i) a random prioritiser as an absolute baseline, and ii) PageRank as an input-naïve prioritiser to assess how predictive centrality is. The comparison of the performance between these four methods was used to assess the predictive power of the label diffusion over three datasets previously introduced and a given network corresponding to an individual pathway database (i.e., KEGG, Reactome, and WikiPathways) or the harmonized integrative network (i.e., PathMe).

As an additional study (see “evaluation by database”), we evaluated whether PathMe, the integrative resource, yields better results than individual pathway databases. To conduct a fair comparison, diffusion algorithms were run on each of the four networks exclusively using entities from the datasets that successfully overlapped with each network. We would like to note that the PathMe network was run using the same input as the individual database it was compared against (i.e., only entities mapped to the individual database were used as an input for PathMe, although the PathMe network could have covered a larger number of entities).

In both evaluations (by method and by database), the input vectors for the propagation were provided as quantitative values for fold-change (in dataset 1) and binarized *p*-value by significance (in dataset 2 and 3). For dataset 1, vectors were codified as the absolute value of the fold change for differentially expressed entities (*p*-value<0.05) while for datasets 2 and 3, these entities were exclusively codified with +1 labels. Subsequently, for each of the three datasets, non-differential entities were given a 0 label and entities not measured remained unlabelled. Normalized scores rely on a permutation analysis that only shuffles labelled nodes. For each entity type, the fold-change (dataset 1) as well as the binarized labels (dataset 2 and 3) were repeatedly split into diffuse-train and validation (50% in each group). Positive labels in diffuse-train were used as an input for the diffusion procedure, whereas those in validation were used to compute the performance metrics. To ensure the most commonly occurring differential entities (e.g. genes) would not dominate over the others, a dedicated diffusion process was carried out stratifying each entity type (section 4.4.3).

After propagation, nodes were prioritised by decreasing scores after calculating their absolute values. The performance metrics used to evaluate the diffusion results were area under the ROC curve (AUROC), as a classical indicator of the overall performance for binary classifiers, and area under the precision-recall curve (AUPRC), as a measure of early retrieval. Finally, Wilcoxon tests were conducted in order to assess the differences in performance between database/method.

We would like to note the computational power required to compute and store the kernel matrix as the main limiting step while running diffusion experiments. This was exacerbated by the large number of experiments conducted to evaluate the robustness of our results (i.e., 100 repeated holdout validations for each input + network + diffusion algorithm combination). As an illustration, the kernel matrix for the PathMe network was nearly 30 GB in size and super computing nodes were required for its calculation (see Subsection 4.5 on hardware). Thus, we have made pre-calculated kernels available for download at https://github.com/multipaths/DiffuPath. Furthermore, our methodology does not take edge directionality or weights into account, which could be addressed in the future through the implementation of alternative diffusion algorithms. Finally, given the organization of the datasets and their heterogeneity, a preprocessing pipeline was implemented to extract the input data (i.e., entity names, fold changes, and/or *p*-values). This step can require a substantial amount of work since one must map and harmonize entity names from the dataset to the network as well as extract and reorganize the relevant information to a proper data structure.

##### 4.4. Results

In this section, we discuss the results of the three validations previously introduced: i) by method, ii) by database, and iii) by entity. All figures presented in this paper and complementary analyses can be reproduced with the Jupyter notebooks located at github.com/multipaths/Results/notebooks/evaluations/repeated_holdout.

The evaluation by diffusion method **(Supplementary Figures 7 to 10)** shows that z normalization yields slightly better performance over raw propagation in dataset 1 using AUROC as a metric, and in dataset 2 and 3 using AUPRC as a metric. However, the Wilcoxon test does not confirm a significant difference when comparing the results of z and raw for dataset 1 for AUPRC, and dataset 2 and 3 for AUROC. When comparing diffusion methods over baselines (i.e., random and PageRank), we can see a significant difference favouring diffusion methods through the range of experiments, thus, validating the predictive power of those approaches on the integrated PathMe network. When comparing the two baselines, PageRank shows slightly better predictive power than random in dataset 2 and 3, suggesting that the differential entities from those datasets tend to be central nodes, while far greater predictive power is achieved by diffusion methods.

The second validation (by database) shows that integrating multiple databases improves performance over individual resources **(Supplementary Figures 11 to 14)**. The results of the validation show that AUROC metrics are higher in the PathMe network for all three datasets. Taking into account the differential mapping of entities for the three individual datasets **(Supplementary Figures 3, 5 and 6)**, DiffuPath improved the coverage over any single-resource network by 40-50% and therefore enhanced the ability of network propagation algorithms to correctly identify genes, metabolites and miRNAs by 10-20%, in terms of AUROC. However, this pattern does not appear when the AUPRC is used as a metric; in these cases, KEGG and Wikipathways outperform PathMe across all three datasets (though Reactome deviates from this trend which may be due to the relatively larger size of the Reactome network in comparison to the KEGG and WikiPathways ones). In such a scenario, the early retrieval performance in smaller networks (e.g. database sub-networks) is more accurate than in larger ones (e.g. integrated PathMe network). This can be attributed to a larger coverage of entities in PathMe which brings down the overall proportion of mapped entities, also leading to lower AUPRCs. For instance, the KEGG dataset includes 6,048 unique HGNC symbols while PathMe contains 13,282 unique HGNC symbols. The number of mapped genes in the first dataset are 1,041 for KEGG and 1,461 for PathMe, which is equivalent to a proportion of 17.2% in KEGG and 11.0% for PathMe, respectively. The lower proportion of mapped entities in PathMe will therefore lower the expected AUPRC.

Finally, we stratified each of the previous experiments by entity to assess the influence of each entity type (i..e, genes, metabolites, and miRNAs) on performance **(Supplementary Figures 15 and 16)**. We found that genes, which represent the largest entity type, were most influential on propagation in all three datasets in terms of AUROC/AUPRC, in contrast to either metabolites or miRNAs. However, because each of these modalities is disproportionately represented and mapped to the network, the predictive power of any one entity type over another remains inconclusive. Nonetheless, it appears that with a greater number of successfully mapped inputs of any entity type, predictive performance is likewise enhanced.

###### 4.4.1. Validation by Method

**Supplementary Figure 7.**
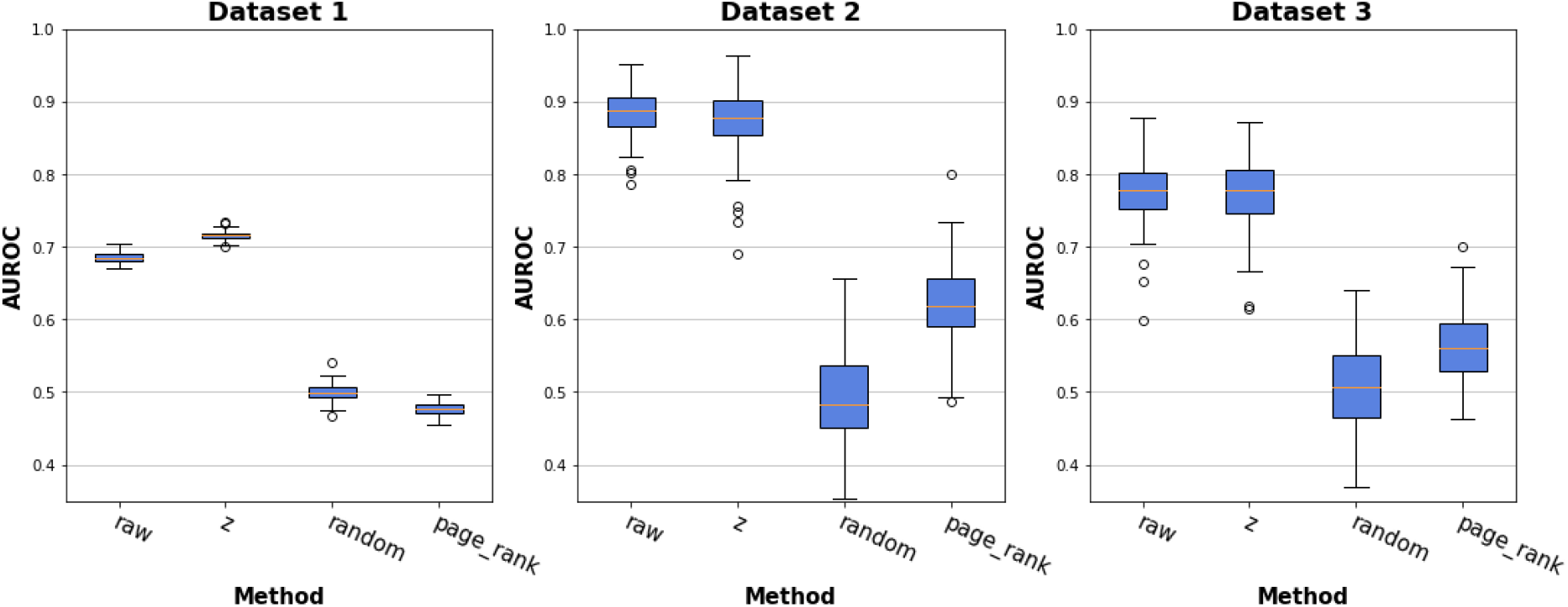
Comparison of prediction performance of diffusion algorithms over the PathMe network using three different multi-omics datasets, and applying different diffusion methods, in order to validate the z-normalization performance over raw diffusion and two baselines, Random and PageRank, this second as a measure of centrality. Each box plot shows the distribution of the AUROC over 100 repeated holdout validations. This figure can be reproduced with the Jupyter notebook located in the git repository at /evaluations/repeated_holdout/by_method.

**Supplementary Figure 8.**
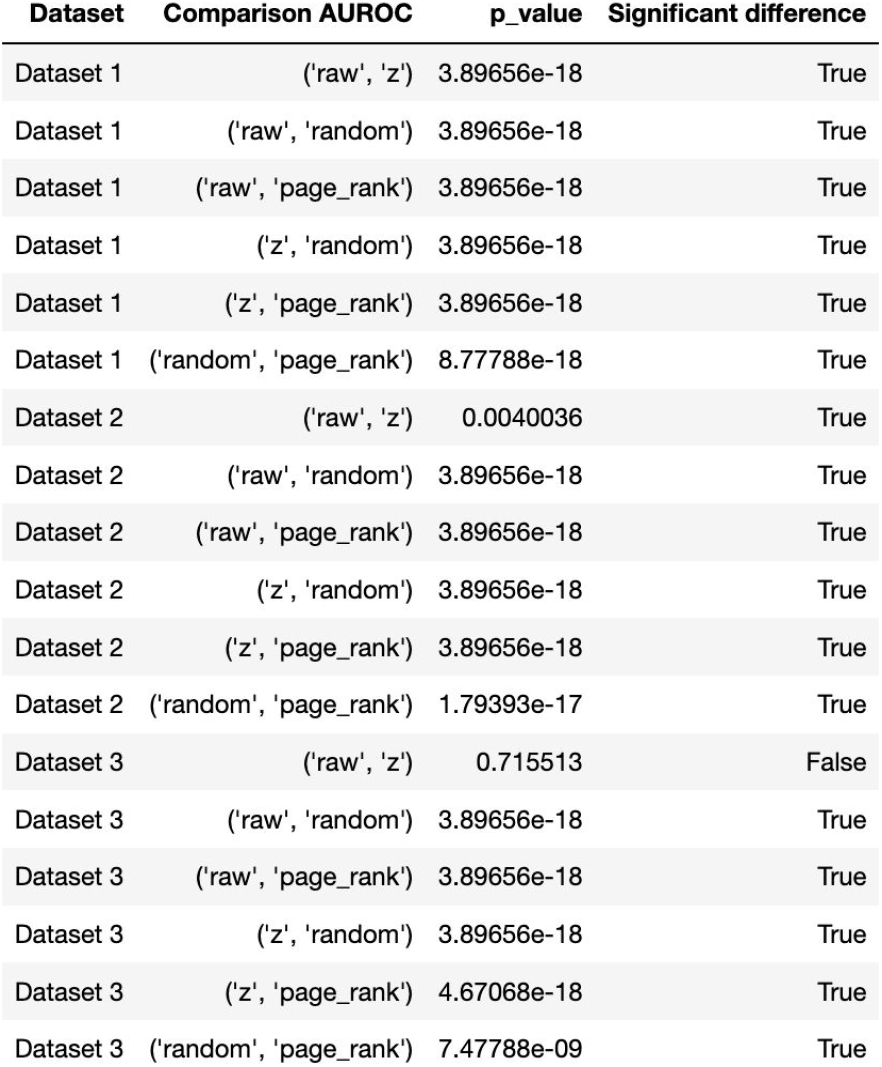
Wilcoxon test to formalize the differential comparisons of prediction performance of diffusion algorithms for the by method validation of AUROC metrics. These results can be reproduced with the Jupyter notebook located in the git repository at /evaluations/repeated_holdout/by_method.

**Supplementary Figure 9.**
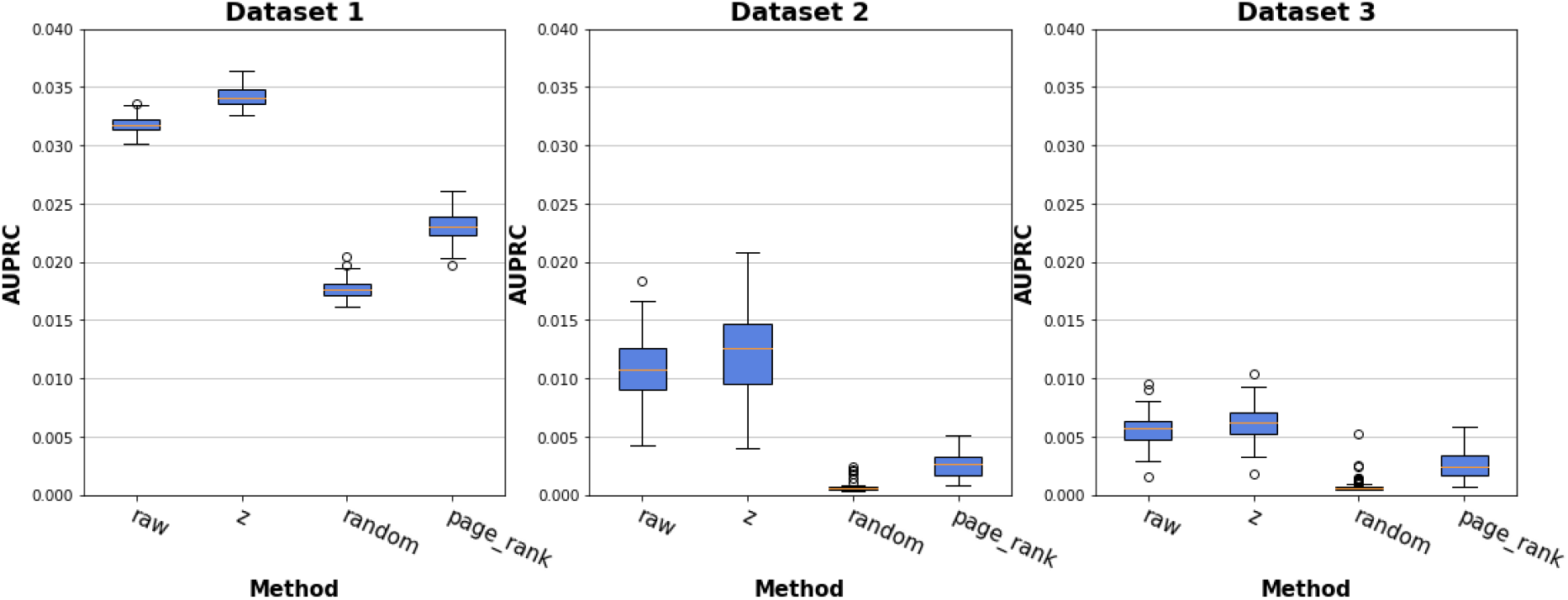
Comparison of prediction performance of diffusion algorithms over PathMe network using three different multi-omics datasets, and applying different diffusion methods, in order to validate the z-normalization performance over the raw diffusion and the two baselines Random and PageRank, the latter of which was used as a centrality measure. Each box plot shows the distribution of the AUPRC over 100 repeated holdout validations. This figure can be reproduced with the Jupyter notebook located in the git repository at /evaluations/repeated_holdout/by_method.

**Supplementary Figure 10.**
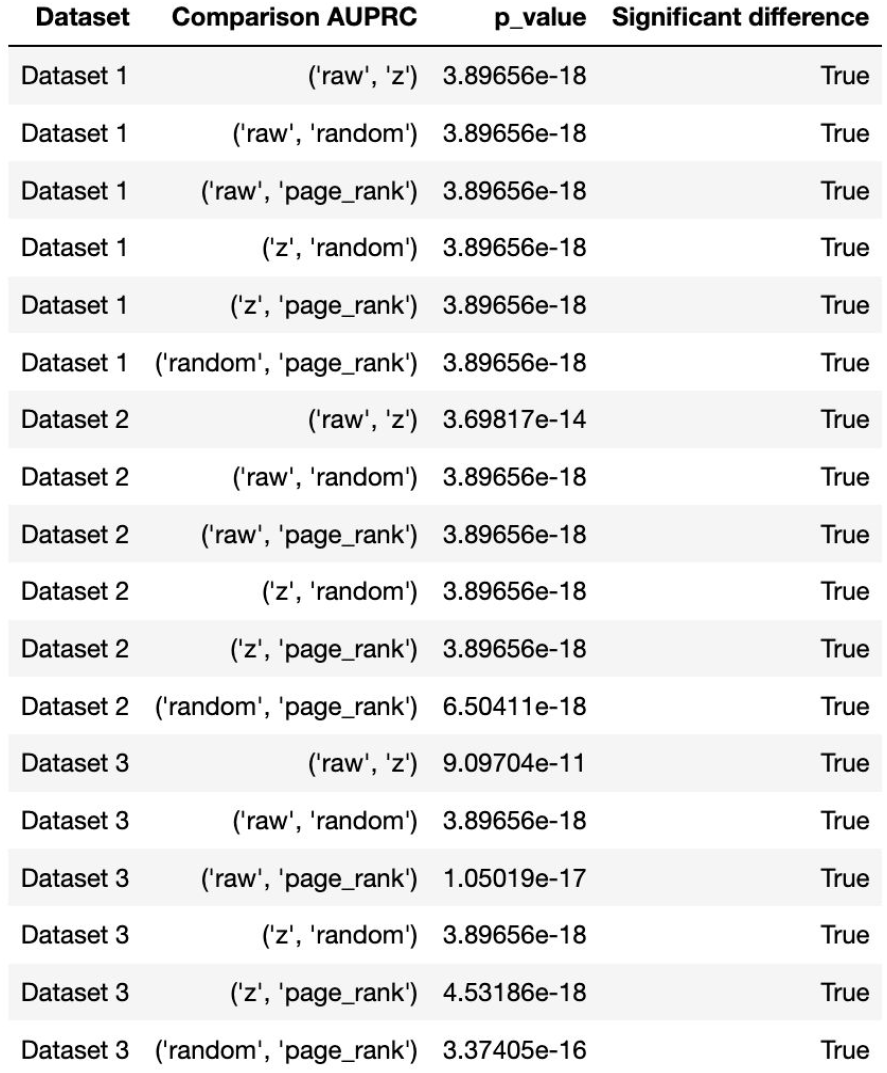
Wilcoxon test to formalize the differential comparisons of prediction performance of diffusion algorithms for the by method validation of AUPRC metrics. These results can be reproduced with the Jupyter notebook located in the git repository at /evaluations/repeated_holdout/by_method.

###### 4.4.2. Validation by Database

**Supplementary Figure 11.**
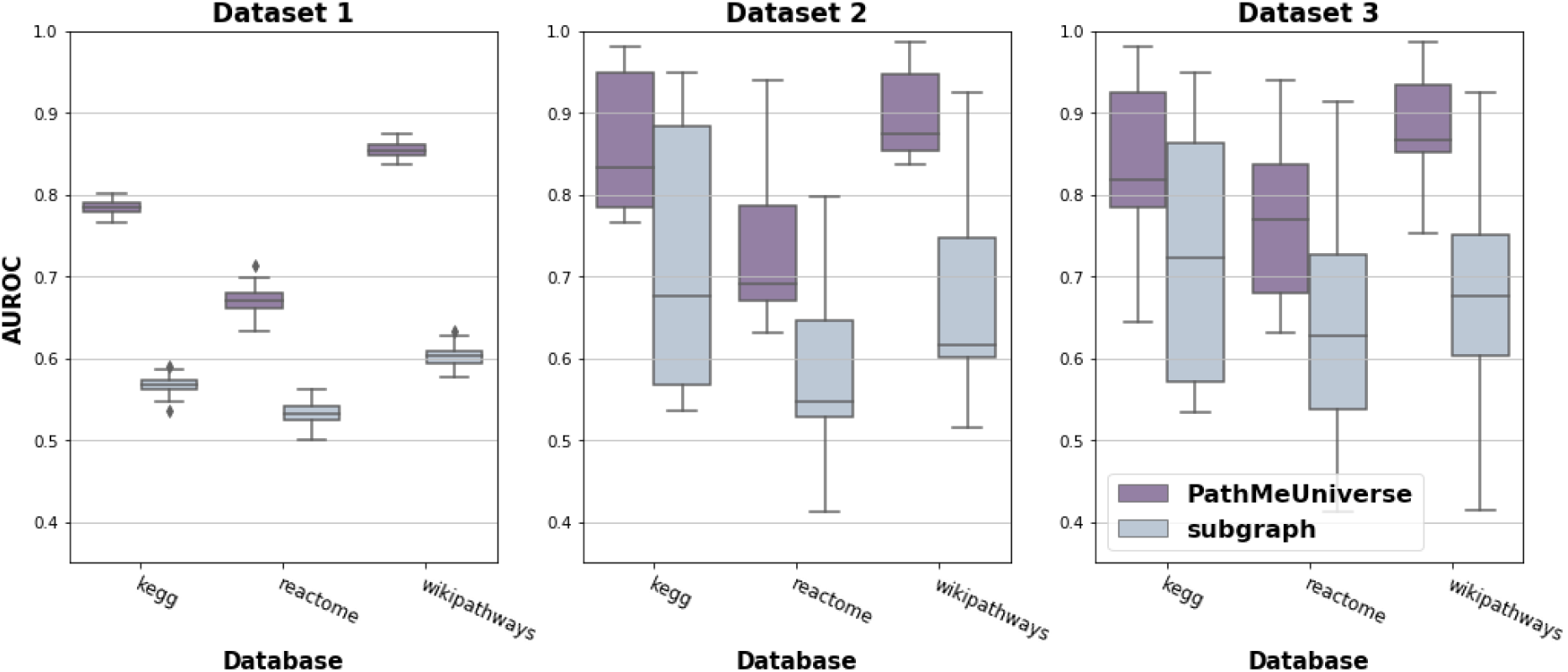
Prediction performance of raw diffusion using three different multi-omics datasets over the integrated PathMe network (purple) and individual pathway databases (blue) in order to validate the PathMe network performance over single-resource pathway databases. Each box plot shows the distribution of the AUROC over 100 repeated holdout validations. This figure can be reproduced with the Jupyter notebook located in the git repository at /evaluations/repeated_holdout/by_database.

**Supplementary Figure 12.**
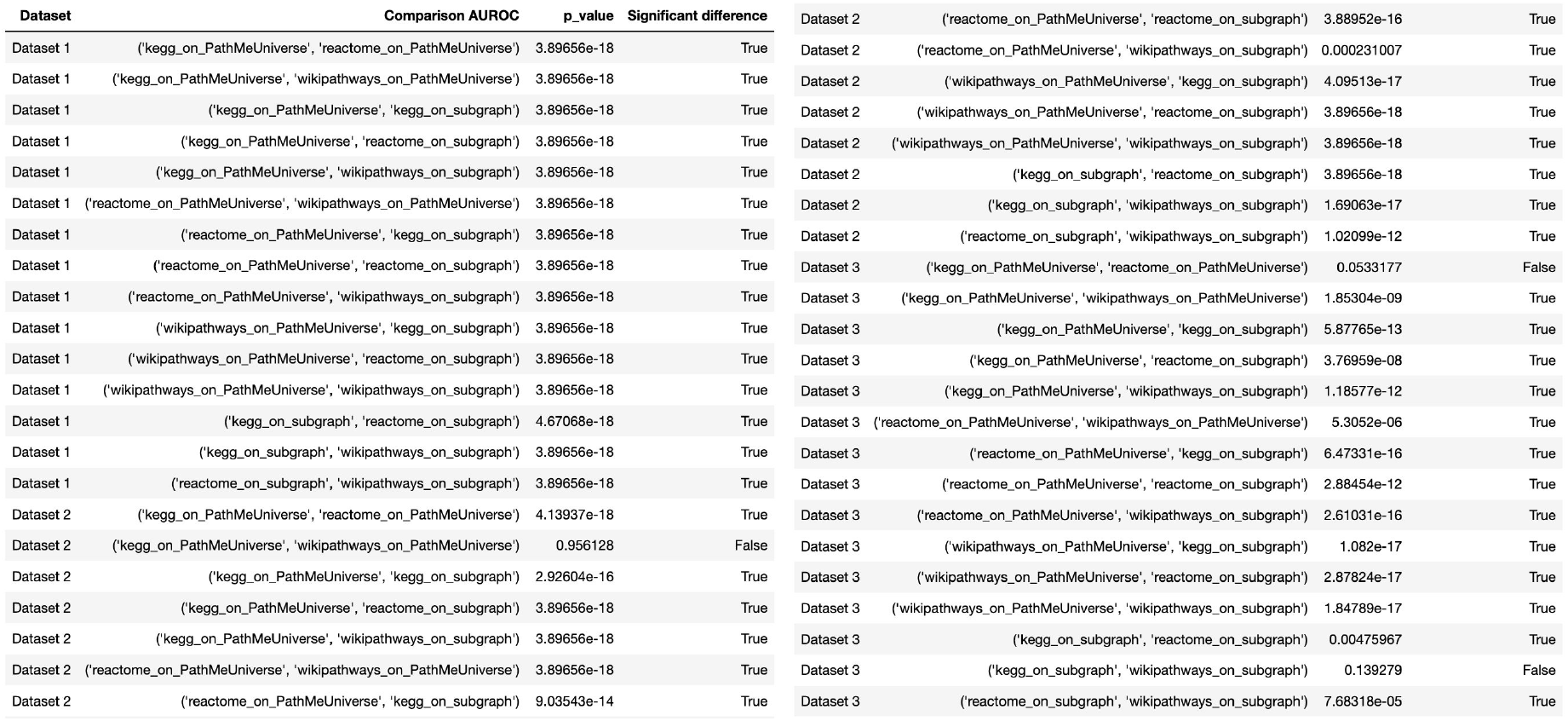
Wilcoxon test to formalize the differential comparisons of prediction performance of diffusion algorithms for the database validation using AUROC as a metric. These results can be reproduced with the Jupyter notebook located in the git repository at /evaluations/repeated_holdout/by_database.

**Supplementary Figure 13.**
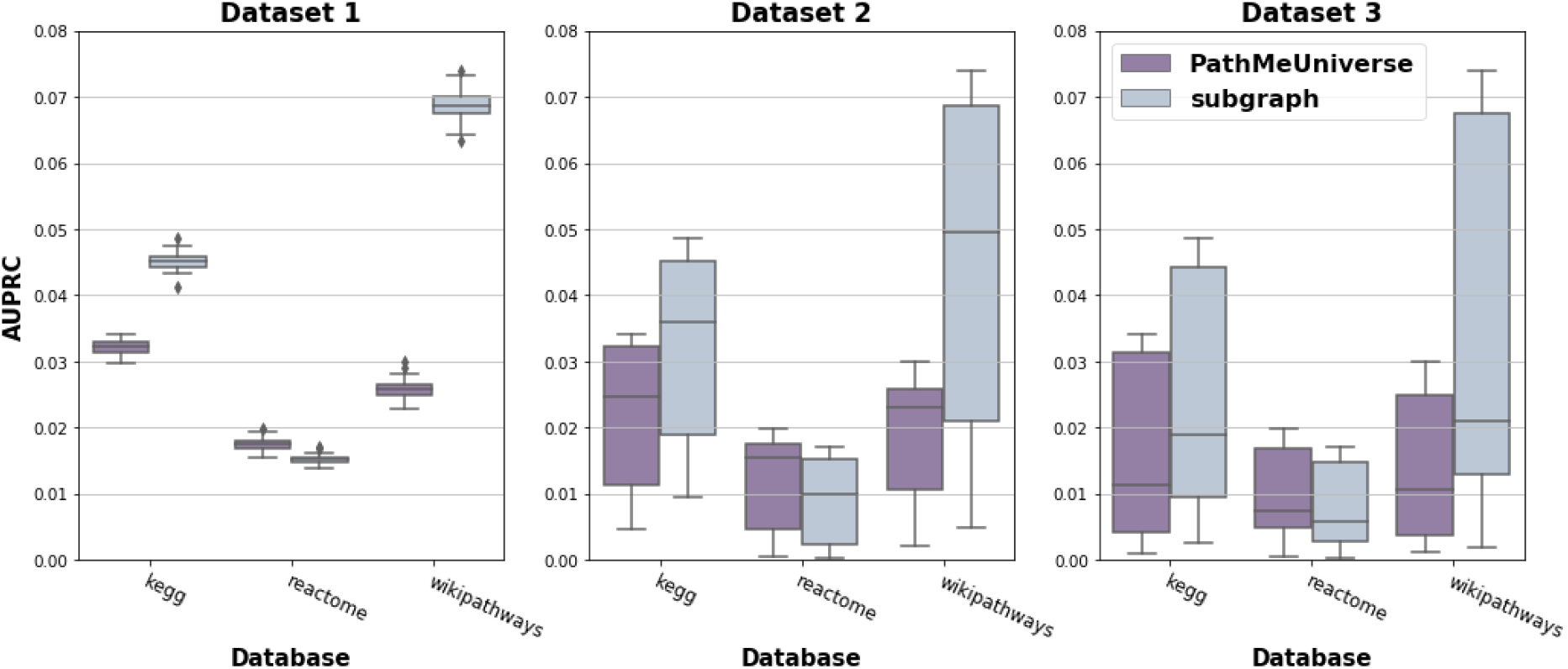
Prediction performance of diffusion using three different multi-omics datasets over the integrated PathMe network (purple) and individual pathway databases (blue) in order to validate the PathMe network performance over single-resource pathway databases. Each box plot shows the distribution of the AUPRC over 100 repeated holdout validations. This figure can be reproduced with the Jupyter notebook located in the git repository at /evaluations/repeated_holdout/by_database.

**Supplementary Figure 14.**
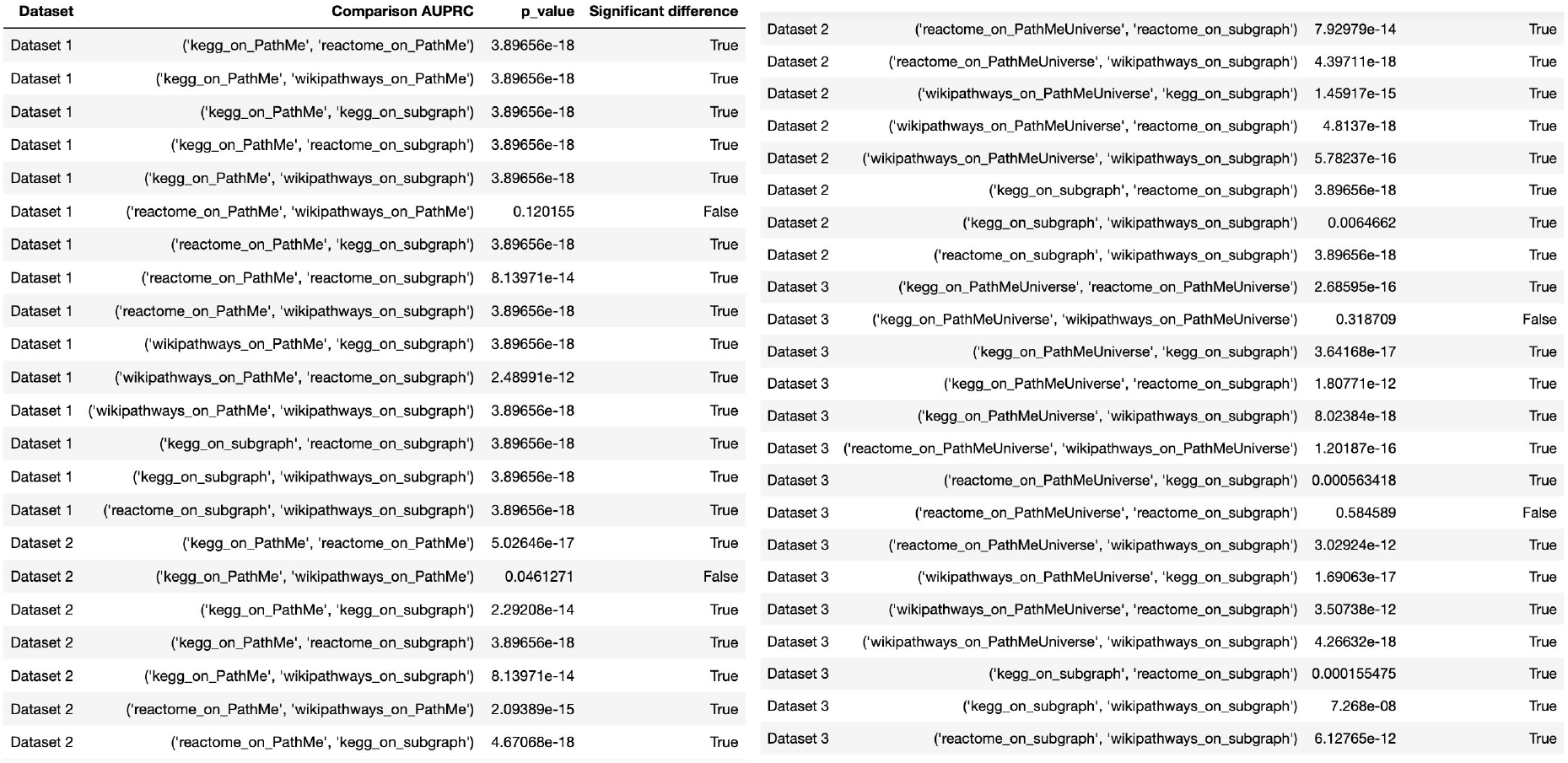
Wilcoxon test to formalize the differential comparisons of prediction performance of diffusion algorithms in the database validation using AUPRC as a metric. These results can be reproduced with the Jupyter notebook located in the git repository at /evaluations/repeated_holdout/by_database.

###### 4.4.3. Validation by method stratified by -omics modality

**Supplementary Figure 15.**
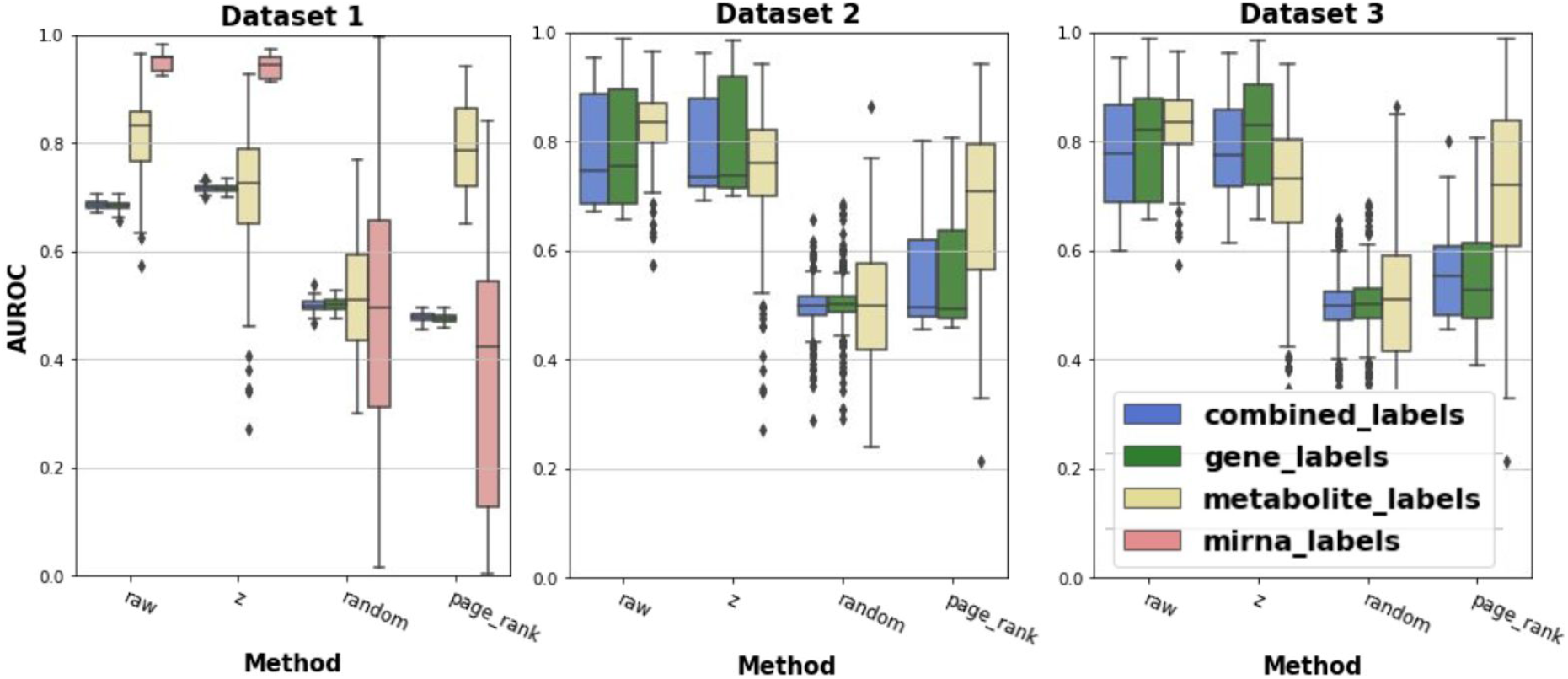
Comparison of prediction performance of diffusion algorithms stratified by entity type over the PathMe network and its corresponding subgraphs. Three different multi-omics datasets and various diffusion methods were applied in order to examine the differential influence of each -omics modality compared to the multi-omics integrated graph. The high variability observed at random and PageRank for miRNAs in dataset 1 can be attributed to the low number of entities in that modality. Each box plot shows the distribution of the AUROC over 100 repeated holdout validations. This figure can be reproduced with the Jupyter notebook located in the git repository at /evaluations/repeated holdout/by_method.

**Supplementary Figure 16.**
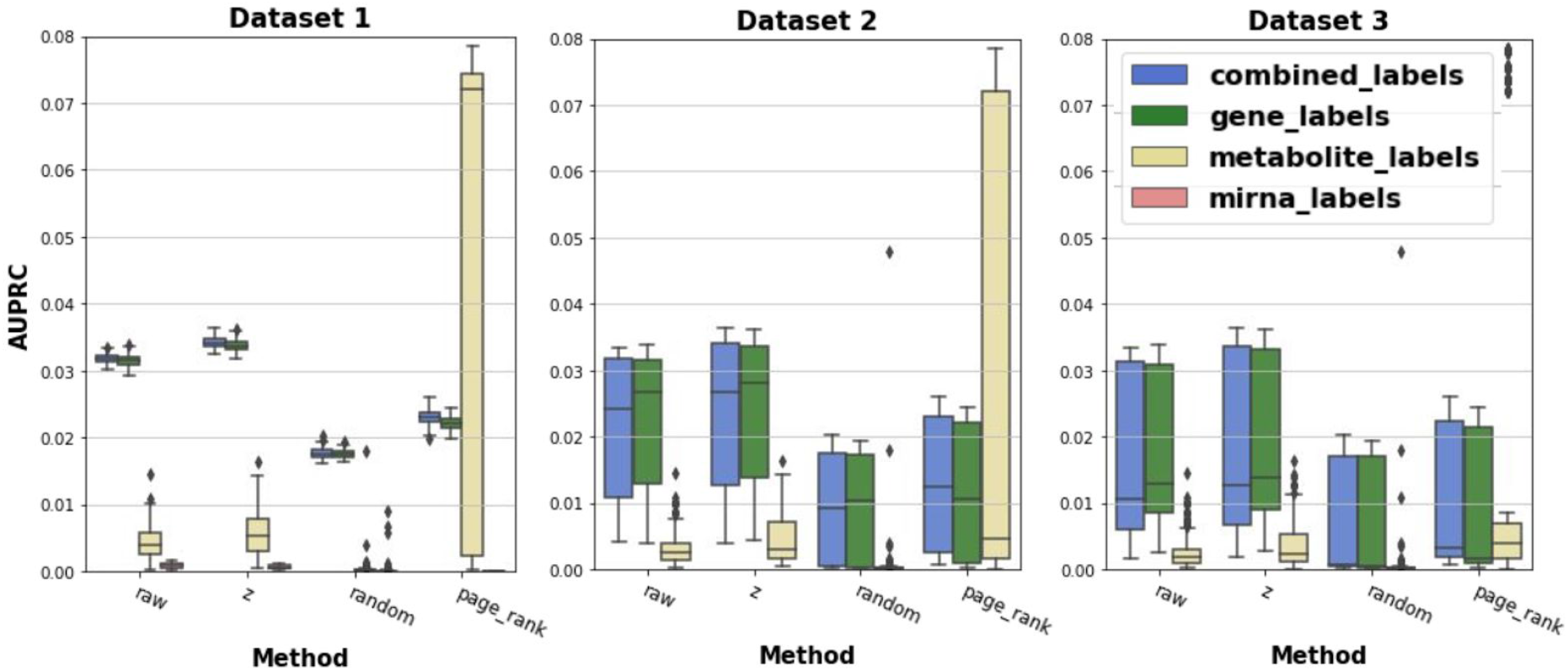
Comparison of prediction performance of diffusion algorithms stratified by entity type over PathMe network using three different multi-omics datasets, and applying different diffusion methods, in order to examine the differential partial influence of each -omics modality compared to the multi-omics integrated graph. Each box plot shows the distribution of the AUPRC over 100 repeated holdout validations. This figure can be reproduced with the Jupyter notebook located in the git repository at [/evaluations/repeated_holdout/by_method](https://github.com/multipaths/Results/tree/master/notebooks/evaluations/repeated_holdout/by_database.ipynb).

##### 4.5. Hardware

Kernel computations were performed on a symmetric multiprocessing (SMP) node with four Intel Xeon Platinum 8160 processors per node with 24 cores/48 threads each (96 cores/192 threads per node in total) and 2.1GHz base / 3.7 GHz Turbo Frequency with 1536GB/1.5TB RAM (DDR4 ECC Reg).

